# National seroprevalence and risk factors of bluetongue virus in domestic ruminants in Peru

**DOI:** 10.1101/2023.07.06.548015

**Authors:** Dennis A. Navarro-Mamani, Jessica Jurado Pucllas, Ana Vargas-Calla, Kevin Ponce Dextre, Yari Zarate Quispe, César A. Murga-Moreno, Perez C Ibelice, Villacaqui A Ruben, Ara G Miguel, Edith Incil, Ortiz O Pedro, Hermelinda Rivera Gerónimo

## Abstract

Bluetongue virus (BTV) is a global infection primarily transmitted among domestic and wild ruminant populations through *Culicoides* spp. midges. Infection is highly prevalent across temperate and tropical regions; however, significant changes in the global distribution of BTV have been observed in recent years. We aimed to evaluate the national BTV seroprevalence and its risk factors among domestic ruminants (cattle, sheep and goats) in Peru. Ruminant sera (cattle= 3 452, sheep= 2 786 and goats=1 568) were collected using a random, two-stage clustering method and analyzed by c-ELISA. The national BTV seroprevalence was 12.7% (95% CI: 11.00-14.63%) and the seropositivity rate for each species was 19.29% (95% CI: 16.0%–31.1%) in cattle, 8.4 (95% CI: 6.6%–10.5%) in sheep, and 9.2% (95% CI: 5.6%–14.8%) in goats. Bivariate and multiple logistic regression analysis were used to evaluate risk factors for BTV seropositivity and to estimate Odds Ratios (*OR*) with 95% CI. We found that age, altitude, and maximum temperature were identified as crucial factors influencing the prevalence of BTV seroprevalence in cattle, sheep and goats. Older animals, particularly cattle and goats between 6 and 24 months and above 24 months, exhibited a significantly higher likelihood of BTV infection. Additionally, higher altitudes above 3000 masl played a protective role, reducing the risk of BTV transmission. Furthermore, increased maximum temperatures, especially exceeding 30 °C, were associated with a greater prevalence of BTV. This study lays the groundwork for identifying BTV serotypes and *Culicoides* spp. in different regions, including altitudes above 3000 masl, to enhance BTV surveillance in Peru.

## Introduction

The global distribution of arboviruses is a severe threat to animal and public health as was evidenced over the last 50 years with the emergence/re-emergence of epidemic arboviral diseases [1]. For livestock, Bluetongue (BT) is an emergent disease with economic importance that affects wild and domestic ruminants worldwide. Bluetongue virus (BTV) is a vector-borne virus transmitted by *Culicoides* spp. between latitudes ∼35°S to 40°–50°N. BTV is endemic where *Culicoides* species and climatic conditions are present in the same geographical space [2,3]. In addition, if vectors migrated to areas outside their normal niche due to climatic and ecological modifications, BT can become epidemic [4–6]. Epidemic bluetongue is estimated to have caused a global economic damage up to US$ 3 billion through decreased animal production, international trade restrictions on live animals and their by-products, and increasing costs of surveillance and control [4,7].

BTV is the type species of the *Orbivirus* genus, belonging to the *Reoviridae* family. BTV has a linear dsRNA genome divided into ten segments that promote segment reassortments and mutations, with the subsequent emergence of new serotypes [8]. To date, more than twenty-eight distinct BTV serotypes have been recognized, each with a different geographic distribution [9,10]. Although *Culicoides* spp. vectors are mainly responsible for BT transmission and infections [2,11,12], horizontal and vertical transmission have also been recorded [13–15]. The severity of BT disease varies based on intrinsic (animal species, breed, age and immune status) and extrinsic factors (presence of competent *Culicoides* and environmental factors) [16]. The acute, clinical course of infection has been reported mainly in sheep, while cattle and goats are usually either asymptomatic or develop only mild symptoms, but with a viremia that can be prolonged for up to 120 days in cattle, making this species a [viral reservoir](https://www.annualreviews.org/doi/abs/10.1146/annurev-virology-110615-042203) [17,18].

Since 1998, Europe has faced sporadic incursions of BTV from other areas and its epidemiology has changed dramatically in recent years, resulting in outbreaks in many countries where it has never been seen before [28,29]. In China, BTV typically infects cattle from intensive farming systems and its presence was also higher in buffalos than dairy cows [30]. Entry of new cattle into herds was noted as another risk factor that was associated with the prevalence of BTV in Brazilian herds [31]. In addition, *Culicoides* spp. are more abundant between 20 and 25 °C, representative of spring and summer seasons, so temperature and precipitation play an important role in BT epidemiology [32]. A number Latin-American countries (Brazil, Argentina, Ecuador, Peru, etc.) have reported BT infection in cattle, sheep, goats and South American camelids (SAC) [19,21] based on serological studies. In Brazil, a 64.81% prevalence has recently been reported in Paraná [22] and 56.44% in Minas Gerais in sheep [23]. Whereas in Ecuador, a 99.1% seroprevalence was recorded in many states from Manabí in cattle [24]. In Peru, a prevalence of BTV higher than 50% has been recorded in sheep from Ucayali [25] and Junin [26], and a 23.8% prevalence in goats from northern Peru [27], However there has been no clinical evidence of BT disease in Peru yet.

Three distinct geographical areas of Peru have been described and of relevance to BT epidemiology: the Pacific coast, the Andean highlands and the Amazonian rainforest [33]. Holstein cattle are principally bred in intensive systems along the Pacific coast, while goats are mostly reared in transhumant systems along the northern coast. In addition, according to the Instituto Nacional de Estadística e Informática (INEI), the largest population of cattle, sheep and CSA are reared in the Andean highlands where people practice either an extensive or semi-extensive farming system. Currently, Peru has not reported any BT outbreak, however serological and molecular evidence of BT was recorded in asymptomatic sheep and cattle from the Amazonian rainforest where *Culicoides insignis* was also recorded, therefore BTV is considered endemic here. Furthermore, according to predictive analytics, *C. insignis* could invade the Andes, where it would find naive populations and potentially cause a massive outbreak in susceptible animals due to variations in surface temperature and rainfall [5,34,35]. The aim of this study was to determine the national seroprevalence and risk factors of bluetongue virus in domestic ruminants (cattle, sheep and goats) of Peru, which would be used for typification studies and surveillance of *Culicoides* spp. vectors in Peru.

## Methods

### Study location

Peru is located at the central-western region of South America, covering an area of 1 285 215.60 km2. It is located between 00°01”48.0” to 04°40”44.5” S and 75°10”29.0” to 81°19”34.5” W [36]. The country has three natural geographic regions with different ecosystems. The Pacific coast (11.7% of the territory) with an altitude of 0 to 500 masl has arid deserts, dry forests, and river basins with productive valleys. Here 44% of the country’s dairy cattle farming is developed, and it is home to 68% of the goat population. The Andean highlands or Sierra (28% of the territory) ranges from 500 to 5500 masl of altitude and has mountainous areas, high plateaus, and inter-Andean valleys with a temperate, cold, or very cold climate [33]. The Andean highlands houses 73%, 94%, 31%, and 100% of the cattle, sheep, goat, and camelid populations, respectively [37]. The Amazon rainforest (57% of the territory) has a tropical and subtropical environment, with an altitude ranging from 500 to 3000 masl, and accounts for about 8% of the cattle population, mainly zebu and their crosses [33].

### Sampling design and serum collection

A two-stage cluster sampling stratified by district was used to obtain a representative sample of 3452 cattle, 2786 sheep, and 1568 goats [38]. Serum sampling were conducted by the Servicio Nacional de Sanidad Agraria del Perú (SENASA) from 2017 to 2019. Sera were transported to the laboratory in accordance with the required conditions. Besides detection of antibodies against BTV, animal-related data [species (cattle, sheep or goat), sex (female or male), and age (< 6 months, 6 to ≤24 months or > 24 months)] and environmental data [altitude (< 1000 masl, 1000 to ≤2 000 masl, 2000 to ≤3000 masl, or > 3000 masl), precipitation (0 to <2 mm/day, 2 to ≤15 mm/day), relative humidity (≤60 %, 60 to ≤80 % or >80 %), and maximum temperature (≤20 °C, 20 to ≤30 °C or >30 °C)] were recoded. No cattle and goat samples from Ancash were collected due to logistical failures. In addition, in the departments of San Martín and Ucayali, no goats were recorded and hence no samples could be collected.

### Detection of antibodies against bluetongue virus

Specific antibodies against the VP7 protein (*Orbivirus* group antigen) of BTV were detected using a commercial competitive ELISA (cELISA) kit (ID Screen® Bluetongue Competition, ID-Vet, Montpellier, France), according to the manufacturer’s instructions. The cELISA kit is widely used in serological studies of BTV globally and is described in the World Organization for Animal Health (formally OIE) manual [39]. The optical density values were measured in a spectrophotometer (Biotek Service & Supplies S.A.) with 450 nm filters and they were calculated for each serum sample as a percentage of competition (S/N%). Serum samples with S/N% values less than 40% were considered positive.

### Statistical analysis

BTV seroprevalence was estimated by calculating the proportion of cELISA-positive animals out of the total number of animals examined, by region and department, with a 95% confidence level. Taylor linearized variance estimators were used due to the staged sampling design [40]. Logistic regression analysis was used to evaluate risk factors for BTV seropositivity and to estimate Odds Ratios (*OR*) with 95% CI. The covariates used in the analysis were altitude, precipitation, relative humidity, and maximum temperature at the farms, and age, sex at the animal level. Initially, bivariate logistic regression was performed to estimate the association of the covariates with seroprevalence. Then, a multiple logistic regression model was formulated, including only the statistically significant variables from the bivariate analyses. All analyses were conducted using the Stata® 17 statistical package (Stata Corp LP, College Station, Texas) with the “svy” command for seroprevalence estimation and logistic regression models from complex sampling. A significance level of 5% was considered.

### Ethics statement

All samples were collected after the agreement of the owners based on “Procedure: collection and sending of samples and requesting tests to the laboratory” with approval number PRO-UCDSA-06 of Unidad del Centro de Diagnóstico de Sanidad Animal. All protocols were approved by the Servicio Nacional de Sanidad Agraria del Perú

## Results

In Peru, the national BTV seroprevalence among ruminants was 12.7% (95% CI: 11.00–14.63%). The BTV seropositivity rate was 19.29% (95% CI: 16.0%–23.1%) in cattle, 8.4 (95% CI: 6.6%–10.5%) in sheep, and 9.2% (95% CI: 5.6%–14.8%) in goats, with regional differences ranging from 0 to 100% (Table 1). No antibodies were detected in ruminants from La Libertad and Tacna and 100% of antibodies were detected in animals from Madre de Dios, Loreto, and Ucayali (Table 1). In general, most of the seropositivity was observed in goat farms from the Northern region, in cattle from the North and Central regions, and in sheep from the South as shown in Figure 1.

**Table 1.** Seroprevalence of the Bluetongue virus at the national, regional and departmental levels in domestic ruminants (cattle, sheep and goats) of Peru

| Region/<br>Departament | Cattle |  |  |  | Sheep |  |  |  | Goat |  |  |  |
| --- | --- | --- | --- | --- | --- | --- | --- | --- | --- | --- | --- | --- |
|  | n | Positive | % | CI 95% | n | Positive | % | CI 95% | n | Positive | % | CI 95% |
| North | 704 | 236 | 33.5 | 26.4 - 41.5 | 417 | 35 | 8.4 | 4.3 - 15.7 | 666 | 127 | 19.1 | 11.1 - 30.9 |
| Tumbes | 34 | 29 | 85.3 | 48.8 - 97.2 | 24 | 9 | 37.5 | 6.8 - 83.1 | 98 | 21 | 21.4 | 8.8 - 43.4 |
| Piura | 137 | 101 | 73.7 | 63.3 - 82.0 | 88 | 24 | 27.3 | 12.9 - 48.6 | 266 | 89 | 33.5 | 14.9 - 59.0 |
| Lambayeque | 29 | 12 | 41.4 | 12.3 - 78.1 | 36 | 2 | 5.6 | 1.0 - 25.2 | 80 | 3 | 3.8 | 0.5 - 23.6 |
| Cajamarca | 366 | 94 | 25.7 | 17.3 - 36.3 | 135 | 0 | 0 | - | 134 | 14 | 10.4 | 5.3 - 19.6 |
| La Libertad | 138 | 0 | 0 | - | 134 | 0 | 0 | - | 88 | 0 | 0 | - |
| Center | 481 | 100 | 20.8 | 13.6 - 30.4 | 874 | 32 | 3.7 | 1.8 - 7.1 | 330 | 0 | 0 | - |
| Ancash* | - | - | - | - | 212 | 2 | 0.9 | 0.1 - 6.9 | - | - | - | - |
| Huánuco | 130 | 57 | 43.9 | 25.3 - 64.2 | 208 | 16 | 7.7 | 1.9 - 26.9 | 84 | 0 | 0 | - |
| Lima-Callao | 121 | 3 | 2.5 | 0.8 - 7.2 | 130 | 0 | 0 | - | 66 | 0 | 0 | - |
| Pasco | 46 | 25 | 54.4 | 23.5 - 82.2 | 112 | 6 | 5.4 | 3.0 - 9.3 | 16 | 0 | 0 | - |
| Junín | 134 | 15 | 11.2 | 4.4 - 25.6 | 184 | 6 | 3.3 | 1.4 - 7.6 | 44 | 0 | 0 | - |
| Ica | 50 | 0 | 0 | - | 28 | 2 | 7.1 | 0.9 - 40.6 | 120 | 0 | 0 | - |
| South | 1862 | 32 | 0.5 | 0.2 - 1.1 | 1408 | 109 | 7.7 | 5.4 - 11.1 | 544 | 0 | 0 | - |
| Huancavelica | 68 | 3 | 4.4 | 0.8 - 20.7 | 176 | 15 | 8.5 | 4.3 - 16.3 | 106 | 0 | 0 | - |
| Ayacucho | 422 | 0 | 0 | - | 176 | 41 | 23.3 | 10.5 - 43.9 | 164 | 0 | 0 | - |
| Apurímac | 294 | 0 | 0 | - | 144 | 5 | 3.5 | 1.2 - 9.4 | 50 | 0 | 0 | - |
| Arequipa | 120 | 0 | 0 | - | 26 | 1 | 3.8 | 0.2 - 48.0 | 76 | 0 | 0 | - |
| Cusco | 228 | 27 | 1.8 | 0.3 - 8.4 | 360 | 23 | 6.4 | 2.9 - 13.6 | 76 | 0 | 0 | - |
| Puno | 696 | 2 | 0.3 | 0.1 - 1.1 | 482 | 22 | 4.6 | 2.6 - 8.0 | 16 | 0 | 0 | - |
| Moquegua | 34 | 0 | 0 | - | 24 | 2 | 8.3 | 0.2 - 77.8 | 20 | 0 | 0 | - |
| Tacna | - | - | - | - | 20 | 0 | 0 | - | 36 | 0 | 0 | - |
| West | 405 | 321 | 79.3 | 66.0 - 88.3 | 87 | 57 | 65.5 | 43.0 - 82.7 | 28 | 17 | 60.7 | 25.2 - 87.6 |
| Amazonas | 80 | 9 | 11.3 | 3.3 - 32.3 | 24 | 2 | 8.3 | 0.5 - 61.7 | 16 | 5 | 31.3 | 8.3 - 69.5 |
| Loreto | 68 | 60 | 88.2 | 73.9 - 95.2 | 8 | 6 | 75 | - | 4 | 4 | 100 | - |
| Madre de Dios | 70 | 70 | 100 | - | 18 | 18 | 100 | - | 8 | 8 | 100 | - |
| San Martín* | 144 | 139 | 96.5 | 77.6 - 99.6 | 12 | 7 | 58.3 | 12.7 - 93.1 | - | - | - | - |
| Ucayali* | 43 | 43 | 100 | - | 25 | 24 | 96 | 69.6 - 99.6 | - | - | - | - |
| Total | 3452 | 666 | 19.3 | 16.0 - 23.1 | 2786 | 233 | 8.4 | 6.6 - 10.5 | 1568 | 144 | 9.2 | 5.6 - 14.8 |

\*In the departments of Ancash, the collection of cattle and goat serum samples could not be carried out due to logistical failures. In addition, in the departments of San Martín and Ucayali, goat samples could not be collected due to the absence of animals.

**Figure 1.**
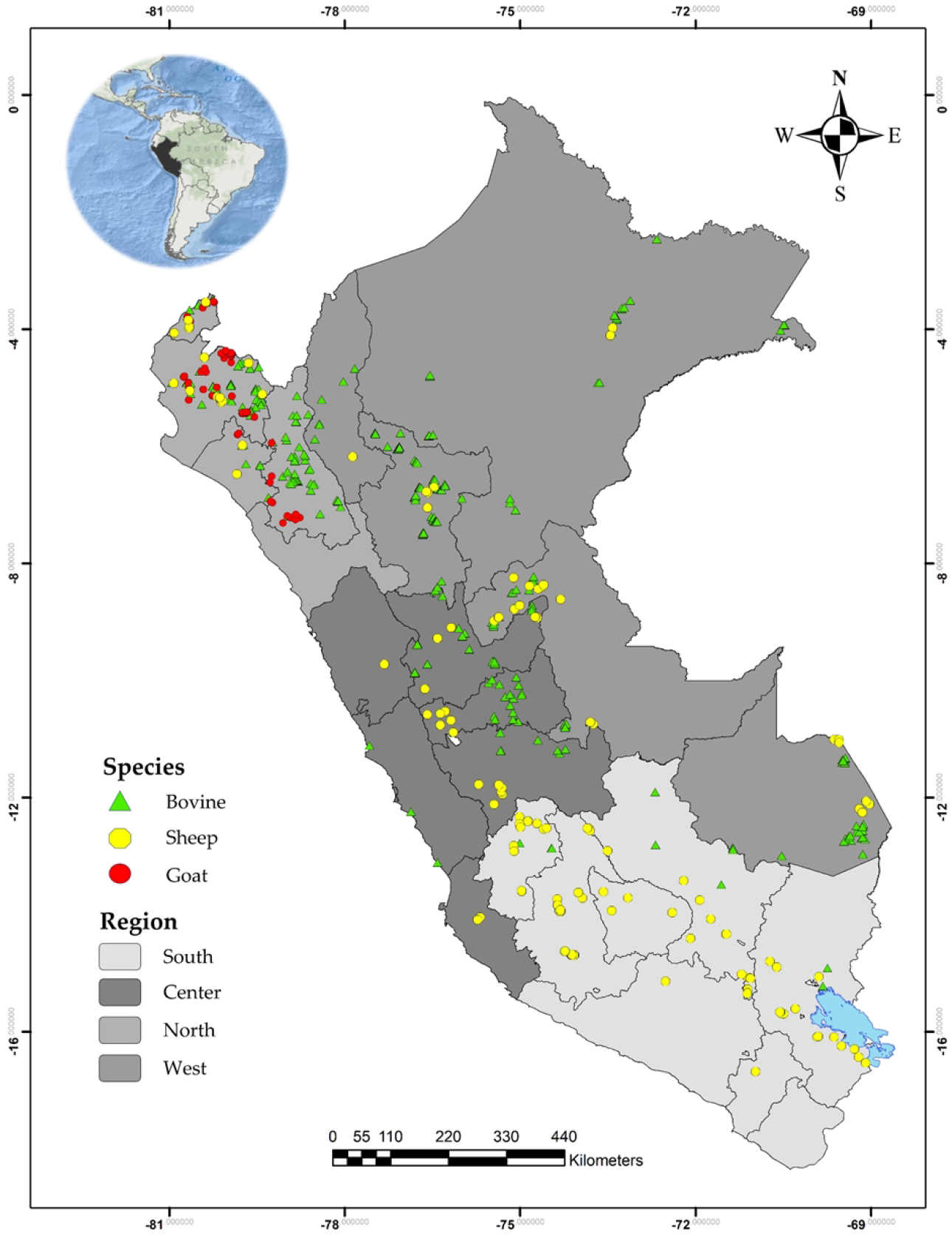
Map of Peru with the departments and region indicating the distribution of farms with BTV seropositive animals. Different gray scale represents different Peruvian region.

### Cattle

Bivariate analysis indicated that increased age, precipitation, relative humidity, and maximum temperature were risk factors for BTV. On the other hand, being male and increasing altitude were protective factors for BTV. The results of multiple logistic regression suggests that at altitudes greater than 3000 masl there was 0.04 (*p* < 0.001) times less likely to have BTV than at altitudes lower than 1000 masl. In addition, we found that in cattle an age older than 24 months (*OR* =1.63, *p* = 0.049), precipitation greater than 2 mm/day (*OR* = 2.67, *p* < 0.001), relative humidity from 60 to 80% and over 80% (*OR* = 3.71, *p* = 0.001 and *OR* = 9.65, *p* < 0.001, respectively), and maximum temperatures greater than 30 °C (*OR* = 7.36, *p* = 0.005) were significantly associated with BTV seroprevalence (Table 2)

**Table 2.** Simple and multiple analysis of cattle showing odds ratios to BTV for different individual and climate characteristics in Peru, 2019

| Variable | Total<br>n | Prevalence<br>% | Simple regression |  |  | Multiple regression |  |  |
| --- | --- | --- | --- | --- | --- | --- | --- | --- |
|  |  |  | OR | CI 95% | p | OR | CI 95% | p |
| Age |  |  |  |  |  |  |  |  |
| ≤ 6 months | 1691 | 12.7 | Ref. |  |  | Ref. |  |  |
| > 6 – ≤ 24 months | 631 | 19.7 | 1.68 | 1.20 – 2.35 | 0.003 | 1.26 | 0.86 – 1.82 | 0.231 |
| > 24 months | 1124 | 29.1 | 2.82 | 1.78 – 4.45 | < 0.001 | 1.63 | 1.00 – 2.67 | 0.049 |
| Sex |  |  |  |  |  |  |  |  |
| Female | 3013 | 20.1 | Ref. |  |  | Ref. |  |  |
| Male | 439 | 14.1 | 0.66 | 0.46 – 0.94 | 0.023 | 0.90 | 0.58 – 1.39 | 0.634 |
| Altitude (masl) |  |  |  |  |  |  |  |  |
| ≤ 1000 | 564 | 62.8 | Ref. |  |  | Ref. |  |  |
| > 1000 – ≤ 2000 | 398 | 46.5 | 0.52 | 0.29 – 0.92 | 0.025 | 1.48 | 0.78 – 2.83 | 0.233 |
| > 2000 – ≤ 3000 | 536 | 18.3 | 0.13 | 0.08 – 0.23 | < 0.001 | 0.91 | 0.39 – 2.16 | 0.833 |
| > 3000 | 1954 | 1.5 | 0.01 | 0.00 – 0.02 | < 0.001 | 0.04 | 0.01 – 0.14 | < 0.001 |
| Precipitation (mm/day) |  |  |  |  |  |  |  |  |
| 0 – ≤ 2 | 1595 | 15.1 | Ref. |  |  | Ref. |  |  |
| > 2 – ≤ 15 | 1857 | 22.9 | 1.68 | 1.09 – 2.59 | 0.019 | 2.67 | 1.56 – 4.59 | < 0.001 |
| Relative humidity (%) |  |  |  |  |  |  |  |  |
| ≤ 60 | 758 | 3.6 | Ref. |  |  | Ref. |  |  |
| > 60 – ≤ 80 | 2220 | 17.5 | 5.73 | 2.78 – 11.85 | < 0.001 | 3.71 | 1.74 – 7.90 | 0.001 |
| > 80 | 474 | 53.0 | 30.47 | 13.50 – 68.80 | < 0.001 | 9.65 | 3.39 – 27.48 | < 0.001 |
| Maximum temperature (°C) |  |  |  |  |  |  |  |  |
| ≤ 20 | 688 | 2.8 | Ref. |  |  | Ref. |  |  |
| > 20 – ≤ 30 | 2177 | 9.0 | 3.48 | 1.24 – 9.80 | 0.018 | 0.64 | 0.20 – 2.07 | 0.456 |
| > 30 | 587 | 76.8 | 116.74 | 40.79 – 334.16 | < 0.001 | 7.36 | 1.84 – 29.42 | 0.005 |

### Sheep

The results of simple logistic regression indicated that latitude was negatively associated, while the maximum temperature was positively associated with BTV seroprevalence. The multiple logistic regression confirmed that temperatures more than 30 °C is a risk factor for BTV (*OR* = 13.34, *p* < 0.001). Also, we founded that sheep between 6 and 24 months were 0.38 (*p* = 0.027) times less likely to have BTV than sheep younger than 6 months and that male sheep were 0.62 (*p* = 0.024) times less likely to have BTV than female sheep (Table 3).

**Table 3.** Simple and multiple analysis of sheep showing odds ratios to BTV for different individual and climate characteristics in Peru, 2019

| Variable | Total<br>n | Prevalence<br>% | Simple regression |  |  | Multiple regression |  |  |
| --- | --- | --- | --- | --- | --- | --- | --- | --- |
|  |  |  | OR | CI 95% | p | OR | CI 95% | p |
| Age |  |  |  |  |  |  |  |  |
| ≤ 6 months | 42 | 11.9 | Ref. |  |  | Ref. |  |  |
| > 6 – ≤ 24 months | 1544 | 7.8 | 0.63 | 0.27 - 1.45 | 0.276 | 0.38 | 0.16 - 0.89 | 0.027 |
| > 24 months | 1198 | 8.9 | 0.72 | 0.30 - 1.71 | 0.451 | 0.59 | 0.24 - 1.48 | 0.260 |
| Sex |  |  |  |  |  |  |  |  |
| Female | 2262 | 8.7 | Ref. |  |  | Ref. |  |  |
| Male | 522 | 7.1 | 0.80 | 0.55 - 1.17 | 0.252 | 0.62 | 0.41 - 0.94 | 0.024 |
| Altitude (masl) |  |  |  |  |  |  |  |  |
| ≤ 1000 | 245 | 33.1 | Ref. |  |  | Ref. |  |  |
| > 1000 – ≤ 2000 | 133 | 15.0 | 0.36 | 0.14 - 0.92 | 0.033 | 0.61 | 0.17 - 2.14 | 0.438 |
| > 2000 – ≤ 3000 | 340 | 6.8 | 0.15 | 0.06 - 0.37 | < 0.001 | 0.68 | 0.17 - 2.66 | 0.577 |
| > 3000 | 2068 | 5.3 | 0.11 | 0.06 - 0.21 | < 0.001 | 0.53 | 0.14 - 2.02 | 0.353 |
| Precipitation (mm/day) |  |  |  |  |  |  |  |  |
| 0 – ≤ 2 | 2235 | 8.7 | Ref. |  |  | Ref. |  |  |
| > 2 – ≤ 15 | 551 | 6.9 | 0.77 | 0.35 - 1.74 | 0.534 | 0.97 | 0.49 - 1.92 | 0.927 |
| Relative humidity (%) |  |  |  |  |  |  |  |  |
| ≤ 60 | 859 | 9.7 | Ref. |  |  | Ref. |  |  |
| > 60 – ≤ 80 | 1663 | 8.0 | 0.81 | 0.47 - 1.40 | 0.454 | 0.65 | 0.35 - 1.20 | 0.166 |
| > 80 | 264 | 6.4 | 0.64 | 0.20 - 2.07 | 0.458 | 0.89 | 0.33 - 2.37 | 0.815 |
| Maximum temperature (°C) |  |  |  |  |  |  |  |  |
| ≤ 20 | 1486 | 4.9 | Ref. |  |  | Ref. |  |  |
| > 20 – ≤ 30 | 1125 | 6.8 | 1.4 | 0.79 - 2.50 | 0.252 | 1.27 | 0.53 - 3.00 | 0.593 |
| > 30 | 175 | 48.0 | 17.87 | 9.33 - 34.21 | < 0.001 | 13.34 | 3.41 - 52.18 | < 0.001 |

### Goats

Bivariate analyze indicated that increasing of age and maximum temperature were risk factors for BTV. While the increase altitude was a protective factor for BTV. The results of multiple logistic regression confirmed that only increasing age was significantly associated with BTV: goats between 6 and 24 months; and goats older than 24 months were 11.19 (*p* < 0.001) and 17.50 (*p* < 0.001) times more likely to have BTV than goats younger than 6 months, respectively (Table 4).

**Table 4.** Simple and multiple analysis of goats showing odds ratios to BTV for different individual and climate characteristics in Peru, 2019

| Variable | Total<br>n | Prevalence<br>% | Simple regression |  |  | Multiple regression |  |  |
| --- | --- | --- | --- | --- | --- | --- | --- | --- |
|  |  |  | OR | CI 95% | p | OR | CI 95% | p |
| Age |  |  |  |  |  |  |  |  |
| ≤ 6 months | 479 | 1.3 | Ref. |  |  | Ref. |  |  |
| > 6 – ≤ 24 months | 839 | 12.5 | 11.28 | 5.01 - 25.41 | < 0.001 | 11.19 | 3.87 - 32.32 | < 0.001 |
| > 24 months | 250 | 13.2 | 11.99 | 5.59 - 25.72 | < 0.001 | 17.50 | 5.64 - 54.30 | < 0.001 |
| Sex |  |  |  |  |  |  |  |  |
| Female | 1526 | 9.3 | Ref. |  |  |  |  |  |
| Male | 42 | 4.8 | 0.49 | 0.11 - 2.13 | 0.338 | 0.51 | 0.11 - 2.42 | 0.390 |
| Altitude (masl) |  |  |  |  |  |  |  |  |
| ≤ 1 000 | 588 | 17.2 | Ref. |  |  | Ref. |  |  |
| > 1 000 – ≤ 2 000 | 230 | 15.7 | 0.89 | 0.51 - 1.58 | 0.699 | 1.57 | 0.83 - 2.95 | 0.163 |
| > 2 000 – ≤ 3 000 | 168 | 4.2 | 0.21 | 0.06 - 0.68 | 0.009 | 0.52 | 0.10 - 2.59 | 0.423 |
| > 3 000 | 582 | 0 | - | - | - | - | - | - |
| Precipitation (mm/day) |  |  |  |  |  |  |  |  |
| 0 – ≤ 2 | 1538 | 9.1 | Ref. |  |  | Ref. |  |  |
| > 2 – ≤ 15 | 30 | 13.3 | 1.54 | 0.30 - 7.87 | 0.603 | 1.58 | 0.18 - 13.83 | 0.679 |
| Relative humidity (%) |  |  |  |  |  |  |  |  |
| ≤ 60 | 468 | 12.8 | Ref. |  |  |  |  |  |
| > 60 – ≤ 80 | 990 | 8.1 | 0.60 | 0.31 - 1.16 | 0.130 | 0.52 | 0.25 - 1.08 | 0.080 |
| > 80 | 110 | 3.6 | 0.26 | 0.05 - 1.28 | 0.097 | 0.46 | 0.05 - 4.54 | 0.506 |
| Maximum temperatura (°C) |  |  |  |  |  |  |  |  |
| > 30 | 462 | 0.4 | Ref. |  |  | Ref. |  |  |
| > 20 – ≤ 30 | 652 | 4.5 | 10.77 | 2.57 - 45.12 | 0.001 | 0.21 | 0.02 - 2.19 | 0.190 |
| ≤ 20 | 454 | 24.9 | 76.66 | 15.94 - 368.74 | < 0.001 | 0.63 | 0.04 - 10.30 | 0.744 |

## Discussion

The current study describes the seroprevalence of bluetongue virus in cattle, sheep, and goats in Peru. This study is particularly important as is the first of its kind to be conducted in the country. The BTV seroprevalence among ruminants was 12.7% (95% CI: 11.00-14.63%). Additionally, the study revealed varying results across different regions and departments, suggesting a possible correlation with the predominant type of animal husbandry practices and the corresponding ecosystems in those areas. For instance, transhumant-type goat farming is the predominant practice in the coastal region characterized by dry forests, and the northern Andean region, characterized by warm to cold climates, provides favorable conditions for livestock development and presence of *Culicoides* spp. [41]. Conversely, the greatest sheep and cattle populations are bred in the central and southern Andes at altitudes about 2 500 to 4 500 masl. The environmental conditions in the central and southern Andean regions are not favorable for the activity of midge vectors that transmit BTV. This might explain the lower BTV seroprevalence observed in these areas compared to the results obtained in the Amazon rainforest region and northern Peru (Table 1 and Fig 1).

The detection of antibodies in animals indicates exposure to BTV in endemic areas. However, the presence of these antibodies in animals from the central and southern highlands may be attributed to several factors, such as the season of the year, climate, precipitation, breeding altitude, management, etc. [42]. These factors may have favored the incursion of midge vectors of the virus from endemic areas of the tropics and subtropics, or the movement of animals. Similar results were reported in populations of yaks from the Tibetan plain in China, where BTV prevalence ranging from 2 to 6% were found at altitudes over 3 000masl because of virus incursions during the short summer season [43,44]. In addition, BTV seroprevalence in Peru was lower than other Latin American countries where BTV is endemic. For instance, cattle showed seroprevalences from 98 to 100% in Ecuador [24,45], dairy cattle from Minas Gerais, Brazil, evidenced 83.3% [46], and cattle from Venezuela around 100% BTV seropositive [47]. In Peru, the Andean region harbors the largest livestock population, characterized by diverse ecological environments and temperatures [36]. While the Amazon rainforest is an endemic area for BTV, factors such as altitude, temperature, and precipitation influence the presence of *Culicoides* species [42]. The Andean Highlands would act as a natural barrier, restricting the migration of *Culicoides* to the coastal and Peruvian Andes.

The study showed age-related effects on BTV antibody seroprevalence in ruminants (Table 2 and 4). Therefore, older cattle and goats (greater than 24 months) have increased odds of being seropositive than younger animals (less < 6 months), as reported by various authors, including Adam *et al*. [48] in cattle, Abera *et al*. [49] in small ruminants, and Bakhshesh *et al*. [50] in cattle, sheep, and goats. This could be associated with the type of breeding, mainly grazing and extensive, and a greater opportunity for contact with midges, particularly during the favorable season of the year [1]. However, sheep’s age was not significantly associated with BTV seroprevalence probably because of their shorter productive life spans (averagely 3-4 years). The results revealed that altitude higher than 3 000 masl was a protective factor (table 2-4). Altitude is undoubtedly a protective factor against BTV infection, possibly limiting *Culicoides* incursions during otherwise favorable conditions, as indicated by some authors [43,44] (Tables 2-4).

Most of sheep are raised in the central and southern regions of the country, specifically in the Andean region (> 3 000 masl). The Andean Highlands provides favorable conditions for sheep farming, including suitable grazing areas, moderate to low temperatures, and access to fresh water sources in a semi-extensive system. We found that altitudes higher than 1000 masl were negatively associated to BTV, and seroprevalence was also low from 3.7% to 7.7% in the center and south respectively (table 1). *Culicoides* midges’ incursions would be associated and potentially certain management activities in sheep, such as shearing. In the central highlands, shearing typically takes place in March, which marks the beginning of autumn, while in the southern regions, it occurs in November or December during the summer season. It is important to note that these periods may coincide with different climatic conditions. In the central highlands, March is generally drier, while in the south, it can be wet. The presence of rainfall can create suitable breeding habitats for *Culicoides* vectors, potentially increasing the risk of BTV transmission during those times [51]. However, further research is needed to investigate the potential impact of sheep management practices on BTV infection rates and transmission risks. Interestingly, antibodies against BTV have been detected in yaks and sheep above 3 000 masl [44,52]. However, information regarding midge vectors of arboviruses that can live at altitudes is scarce. In Colombia, several species of *Culicoides* have been identified at altitudes around 2 500 masl [53]. Additionally, in recent decades, global warming may facilitate the migration of *Culicoides* species from warmer ecosystems to higher altitudes, where susceptible animal populations can be found.

Our study also revealed that temperatures above 30 °C were identified as a risk factor for BTV seroprevalence (Tables 2-4). It is evident that temperatures ranging from 20 to 25 °C during spring and summer, along with factors such as precipitation, wind speed, and rainfall, influenced the abundance of *Culicoides*, which are associated with BT infection [12,54]. The Amazonian rainforest, located within the Neotropic and the Amazon basin, comprises tropical and subtropical moist broadleaf forests [55], providing an endemic environment for BTV due to the presence of its *Culicoides* vector [1]. The favorable temperature conditions in this region facilitate rapid BTV replication, shorten the intervals between *Culicoides* feeds, and increase the risk of transmission. Additionally, higher temperatures accelerate the development of *Culicoides* eggs and larvae, while potentially impacting the lifespan of female *Culicoides* [12,56,57].

Interesting studies on global warming and climate change indicate that their effects will have a significant impact on public health and animal health in different regions of South America and Peru [34,35,58]. Peru is one of the countries most vulnerable to climate change, and certain events such as El Niño–Southern Oscillation (ENSO) phenomenon could become more intense, creating favorable conditions for the distribution and survival of *Culicoides* vectors of BTV in the Andean region, especially in the north. While clinical cases of BTV disease have not been reported to date, it is necessary to identify the BTV serotypes, as well as the species of the *Culicoides* genus, including *C. insignis*, present in different ecosystems of the country. These data are essential for active serological and molecular surveillance by the Peruvian health authorities.

## Conclusions

This study provides basic epidemiological information on Bluetongue virus (BTV) in domestic ruminants in Peru. BTV is widespread nationwide with a low seroprevalence in cattle, sheep and goats, but it is endemic in the rainforest amazon. BTV seroprevalence moderately differs based on the population size of ruminants in different regions, with goats predominant in the northern regions, cattle in the central and southern regions, and sheep mainly in the southern region of the country. Risk factors associated with BTV infection varied among the evaluated ruminant species and their respective regions of origin. The study serves as a foundation for further identifying circulating BTV serotypes nationwide and identifying *Culicoides* spp. in different regions, including altitudes above 3 000 masl, as a protective factor for enhancing active surveillance of BTV in Peru’s major livestock-populated areas. Although there is currently no evidence of clinical disease in ruminants in the country, it could potentially occur in the near future due to climate change.

